# Laser Recording of Subcellular Neuron Activities

**DOI:** 10.1101/584938

**Authors:** Yu-Cheng Chen, Xuzhou Li, Hongbo Zhu, Wei-Hung Weng, Xiaotian Tan, Qiushu Chen, Xueding Wang, Xudong Fan

## Abstract

Advances in imaging and recording of neural activities with a single neuron resolution have played a significant role in understanding neurological diseases in the past decade. Conventional methods relying on patch-clamp and electrodes are regarded as invasive, whereas fluorescence-based imaging tools are useful but still suffer from a low signal-to-noise ratio and low sensitivity. Here we developed a novel optical imaging and recording system by employing laser emissions to record the action potentials in single neurons and neuronal networks caused by subtle transients (Ca^2+^ concentration) in primary neurons *in vitro* with a subcellular and single-spike resolution. By recording the laser emissions from neurons, we discovered that lasing emissions could be biologically modulated by intracellular activities and extracellular stimulation with >100-fold improvement in detection sensitivity over traditional fluorescence-based measurement. Finally, we showed that ultrasound can wirelessly activate neurons adsorbed with piezoelectric BaTiO_3_ nanoparticles, in which the neuron laser emissions were modulated by ultrasound. Our findings show that ultrasound stimulation can significantly increase the lasing intensity and neuron network response. This work not only opens the door to laser emission recording of intracellular dynamics in neuronal networks but may provide an ultra-sensitive detection method for brain-on-chip applications, optogenetics, and neuro-analysis.

---

Intracellular recording of action potentials in cultured neurons is critical to studying neural activities and preclinical neurological diseases^1-4^. Both spontaneous and stimulated spiking activities provide unique information on the interactions of ion channels and synapses from single neurons to complex networks^5,6^. For decades, a number of techniques have been developed to record action potentials, including patch-clamps^7,8^, optical-probes^3,9,10^, multi-electrode arrays^6,11,12^, and several types of optical imaging modalities^4,5,10,11^. In particular, single/two-photon fluorescence microscopy has become one of the most powerful and popular approaches to monitoring ion dynamics of neuronal activities in large networks. However, fluorescence-based measurements are not adequately sensitive to detect or monitor the subtle changes in ion concentrations (e.g., Ca^2+^), as the relative fluorescence change (ΔF/F) for a single action potential is usually on the order of only 0.2-0.5^2,13^. Consequently, minute changes in spontaneous activities may often be missed as noise. The low sensitivity and low signal-to-noise (SNR) of fluorescence imaging also result in deteriorated temporal and spatial resolution, both of which require heavy post-signal processing. Therefore, how to improve the signal quality of optical recording during neuronal spiking activity *in vitro* remains challenging.

To date, laser emission from biological materials (also known as *biolasers*^14-21^) has demonstrated distinct advantages over fluorescence in terms of signal amplification, narrow linewidth, and strong intensity, as well as the ability to eliminate unwanted fluorescence or scattering background, leading to orders of magnitude increase in detection sensitivity^22-24^, SNR^25,26^, and signal-to-background ratio^25,27^. Recently, a plethora of researches have shown the promise of cellular lasers, including cell tracking^28-30^, labeling^30,31^, and monitoring^24^. However, the ability of the biolaser in highly sensitive intracellular sensing with high spatial and temporal resolution has not yet been fully exploited.

In this article, we demonstrated a “neuron laser” and subsequently employed laser emission microscopy to record the action potentials in single neurons and neuronal networks caused by subtle transients (Ca^2+^ concentration, nM level) in primary neurons *in vitro* with a subcellular and single-spike resolution. For the first time, we showed that lasing emissions could be biologically modulated by intracellular activities and extracellular stimulation of neurons and that laser emission-based recording of spontaneous neuron spiking has >100-fold improvement in sensitivity to detect intracellular calcium dynamics, compared to conventional fluorescence-based recording. Moreover, by using a low-intensity pulsed ultrasound to activate neurons absorbed with piezoelectric BaTiO_3_ nanoparticles, we were able to record stimulated neuron spiking with the neuron laser, which shows even higher lasing intensity changes (or sensing signals) and increased network responses in the presence of acoustoelectric stimulation. Our work will lead to a technology for better visualization of spatial/temporal neural transients that may reveal valuable activities governing electrophysiological processes in neurobiology.

Fig. 1(a) illustrates the concept of a neuron laser. Primary cortical neurons labeled with a calcium indicator, such as Oregon Green BAPTA-1 (OGB-1), serve as the laser gain medium, whereas a Fabry-Pérot (FP) microcavity is formed by two highly-reflective dielectric mirrors. The inset shows the fluorescence image of neurons stained with OGB-1 cultured on the bottom mirror. Both spontaneous and stimulated neuron activities were recorded by imaging the laser emission from neurons with a CCD camera. The unique threshold behavior of the neuron laser can be explored for sensitive detection of spontaneous neuronal activities, as illustrated in Fig. 1(b). At the resting potential, no neuronal activity and hence no Ca^2+^ spiking occurs. Therefore, the laser is operated below its lasing threshold and only extremely weak fluorescence background was observed. When a spontaneous/stimulated spiking occurs, the increased Ca^2+^ concentration boosts the neuron laser to operate above its lasing threshold, resulting in the emergence of lasing emission. This sharp cut-off characteristic of the neuron laser below and above the lasing threshold ensures orders of magnitude improvement in detection sensitivity.

**Figure 1.**
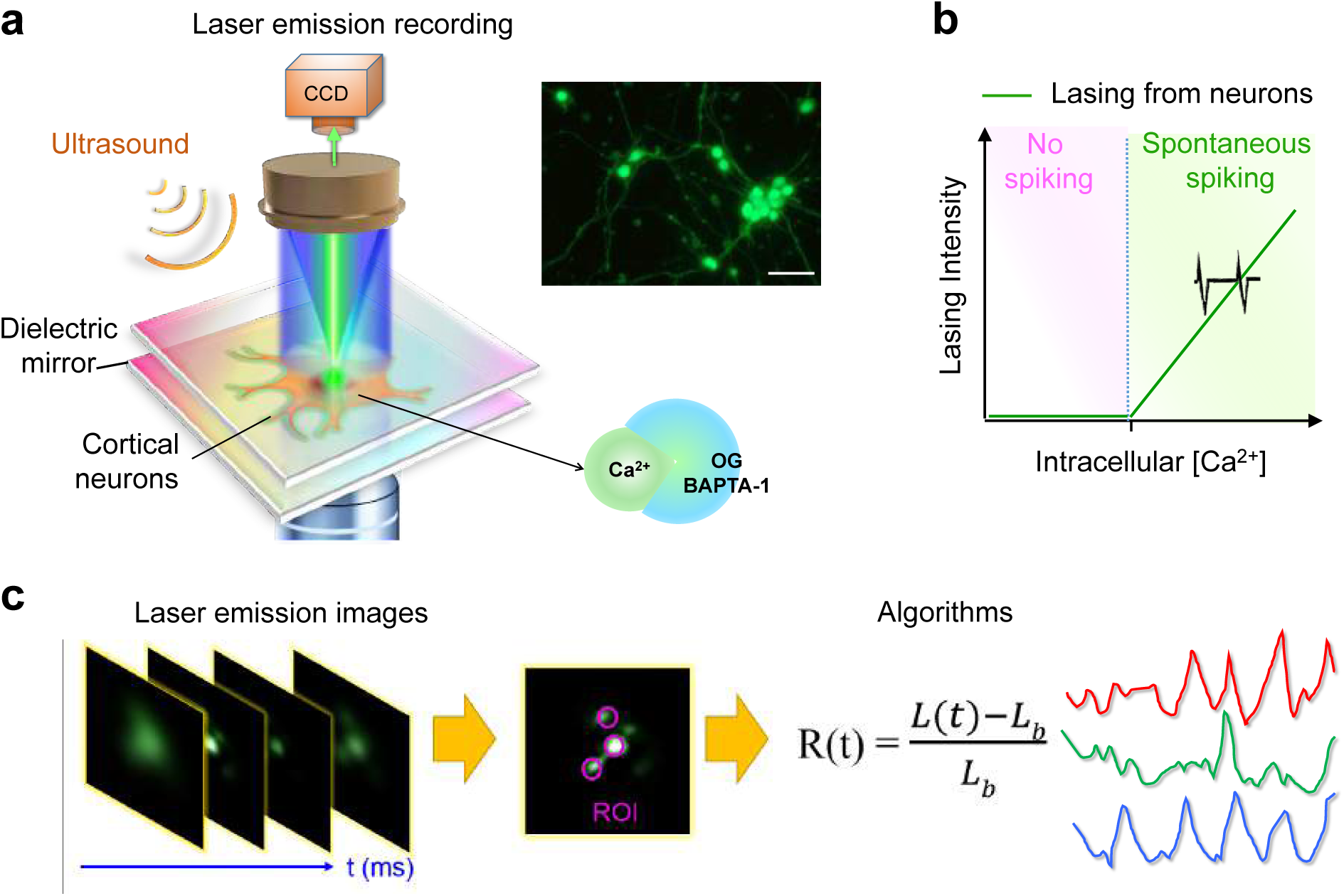
Concept of optical recording of neuron spikings with a neuron laser. **(a)** Schematic of the experimental configuration of a neuron laser, in which neurons are sandwiched inside an FP cavity formed by two highly reflective mirrors. Both spontaneous and ultrasound stimulated neuron spiking activities are recorded with laser emission. The inset shows a fluorescence image of OGB-1 labeled primary rat cortical neurons cultured on mirrors. Scale bar, 50 µm. **(b)** Output intensity from neuron lasers as a function of intracellular Ca^2+^ concentration. Under the fixed pump intensity, only when intracellular Ca ^2+^ reaches above the baseline concentration (∼50 nM) caused by spontaneous or stimulated spiking can laser emission emerge. **(c)** Protocol of image processing to monitor neuronal activities. Different regions of interest (ROI) are selected on the laser emission images over a period of time, then calculated by the algorithms developed in-house.

Optical recording of neurons was performed by collecting series of laser emission images from a CCD camera, as illustrated in Fig. 1(c). The recorded laser images over time were then transformed by the equation:

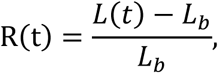

where L(t) is defined as the measured laser emission intensity at time, t, from a certain region of interest (ROI) on the laser emission microscopic image. L_b_ is defined as the minimum laser emission intensity within a period of time, which is used as the baseline signal. More specifically, L(t) is the laser emission during a calcium transient (or action potential), whereas L_b_ is the “emission” when there is no calcium transient. R(t) is the relative change of the laser emission intensity at time, t. Note that the definition of R(t) is analogous to the relative change of the fluorescence signal, ΔF/F, used in fluorescence microscopy.

## Results

### Lasing with calcium indicators

As a control experiment, we first used OGB-1 to demonstrate laser-based detection of subtle changes of free Ca^2+^ concentration by using the same setup in Fig. 1(a). OGB-1 is well known as a calcium indicator that has an increased fluorescence upon the binding to free Ca^2+^. The fluorescence emission spectra of OGB-1 in the presence and absence of Ca^2+^ are shown in Fig. 2(a), in which the relative fluorescence change (ΔF/F) is less than 0.1. For comparison, the laser emission spectra of OGB-1 (pure dye) and OGB-1 with Ca^2+^ are plotted in Fig. 2(b). We can clearly see that no laser emission could be observed from pure OGB-1; however, a huge laser emission emerged when OGB-1 was mixed with Ca^2+^. In contrast to Fig. 2(a), the relative laser intensity change, R(t), is more than 100, 3 orders magnitude larger than that in conventional fluorescence. In later sections, we will show how to translate the same laser emission based detection technology to sensitively monitor the subtle intracellular activities such as spontaneous/stimulated spiking in living neuron cells.

**Figure 2.**
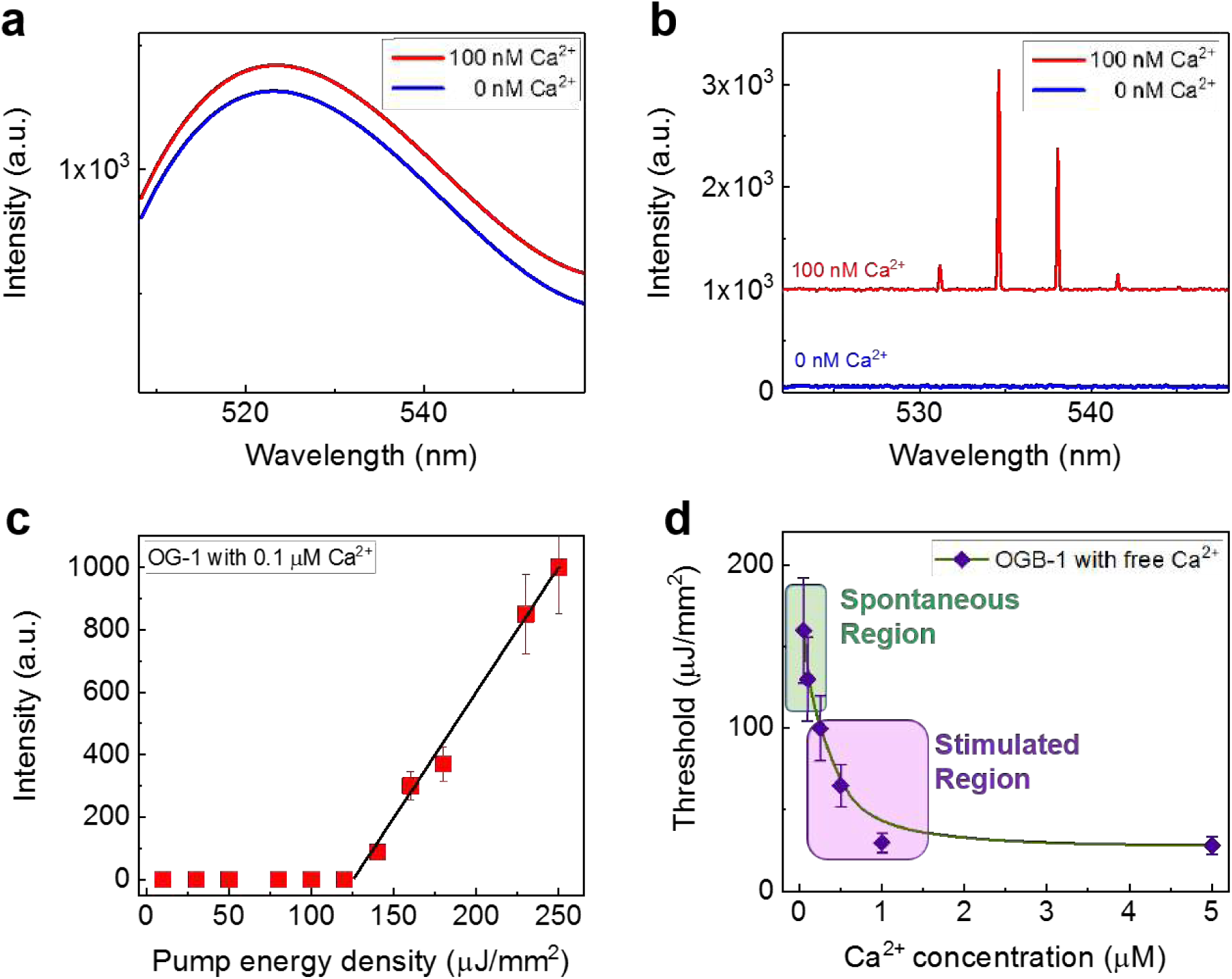
Lasing with calcium indicator OGB-1. **(a)** Fluorescence emission spectra of pure OGB-1 (blue curve) and OGB-1 mixed with 100 nM of free Ca^2+^ (red curve). **(b)** Laser emission spectra of pure OGB-1 (blue curve) and OGB-1 mixed with 100 nM free Ca^2+^ (red curve) under a pump energy density of 280 μJ/mm^2^. The lasing emission curve is vertically shifted for clarity. **(c)** Spectrally integrated (530 nm – 545 nm) laser output as a function of pump energy density extracted from the spectra in (b). The solid line is the linear fit above the lasing threshold, showing a lasing threshold of 120 μJ/mm^2^. Error bars are obtained based on the power variation of the pump laser. **(d)** Lasing threshold calibration of pure dye (OGB-1) mixed with different concentrations of free Ca^2+^. The shaded green area depicts the spontaneous calcium spiking region (50-200 nM). The shaded pink area depicts the stimulated calcium spiking region (150-1500 nM). Excitation wavelength: 475 nm.

The corresponding spectrally integrated laser emission versus pump energy density presented in Fig. 2(c) shows the lasing threshold of approximate 120 μJ/mm^2^. To characterize the OGB-1 laser, in Fig. 2(d) we investigated the lasing threshold of pure OGB-1 (50 μM) mixed with various concentrations of free Ca^2+^. Note that in Fig. 2 we used this concentration to mimic that in neurons because the effective final intracellular OGB-1 concentration in neurons is measured to be 50-60 μM using our staining process. It is shown that the lasing threshold increases with the decreased Ca^2+^ concentration, which is expected. In particular, a drastic difference in the lasing threshold can be seen when the Ca^2+^ concentration is in the range of 50 nM to 100 nM, which suggests that OBG-1 and its lasing emission can be used to monitor the subtle transients of spontaneous calcium spike activities, as the spontaneous Ca^2+^ concentration in neurons is within the same range of 50 nM to 100 nM. As for stimulated calcium spike activities, a higher Ca^2+^ concentration ranging from 100 nM to 1500 nM is expected.

### Lasing in living neurons

Next, in Fig. 3(a) we demonstrated a “neuron laser” when we used 25 μM OGB-1 solution to label the neuron *in vitro* (note that the actual OGB-1 concentration inside a neuron cell was measured to be 50 – 60 μM). A single mode lasing profile was first observed when the pump energy density is slightly above the lasing threshold. Additional lasing modes emerge as the pump increases. The signal-to-noise ratio (SNR) defined as the ratio between the lasing peak intensity and the background noise away from the lasing peak wavelength) is over 500. The inset shows an example of a bright field image of a single neuron with laser emissions within the cell soma. The corresponding spectrally integrated laser emission versus pump energy density presented in Fig. 3(b) shows the lasing threshold of approximate 60 μJ/mm^2^ for the neuron at the resting potential. As visualized by the CCD images in the inset of Fig. 3(b), no lasing is observed when the pump was below 60 μJ/mm^2^. In contrast, sharp lasing emission emerges as the pump intensity increases drastically. Subcellular laser emissions can be observed only from specific sites in the neuron. Note that although the neuron in Fig. 3(a) was at the resting potential (no spontaneous Ca^2+^ spiking), the lasing emission could still be obtained, but at relatively high pump energy densities. In contrast, the lasing threshold becomes much lower when the neuron undergoes a spiking process. In most cases, only a single lasing spot with a single lasing mode is observed for each neuron under low pump intensities. When the pump intensity is increased, multiple lasing spots can also be observed within a single neuron as exemplified in Fig. 3(c), in which each spot represents one individual laser and has its own lasing threshold due possibly to slightly different local dye concentration in the neuron cell. This phenomenon can be used in the future to characterize the intracellular dye concentration. Note that only at very high pump intensities (a few times the threshold) can multiple lasing sites be observed within a single neuron. Therefore, it is not difficult for us to tell whether the multiple lasing spots are from a single neuron or multiple neurons. Finally, in Fig. 3(d) we investigated the spatial resolution of such a neuron laser. The inset shows that lasing emission could be achieved from both the soma and dendrites of neuron, and the FWHM was measured to be approximately 950 nm.

**Figure 3.**
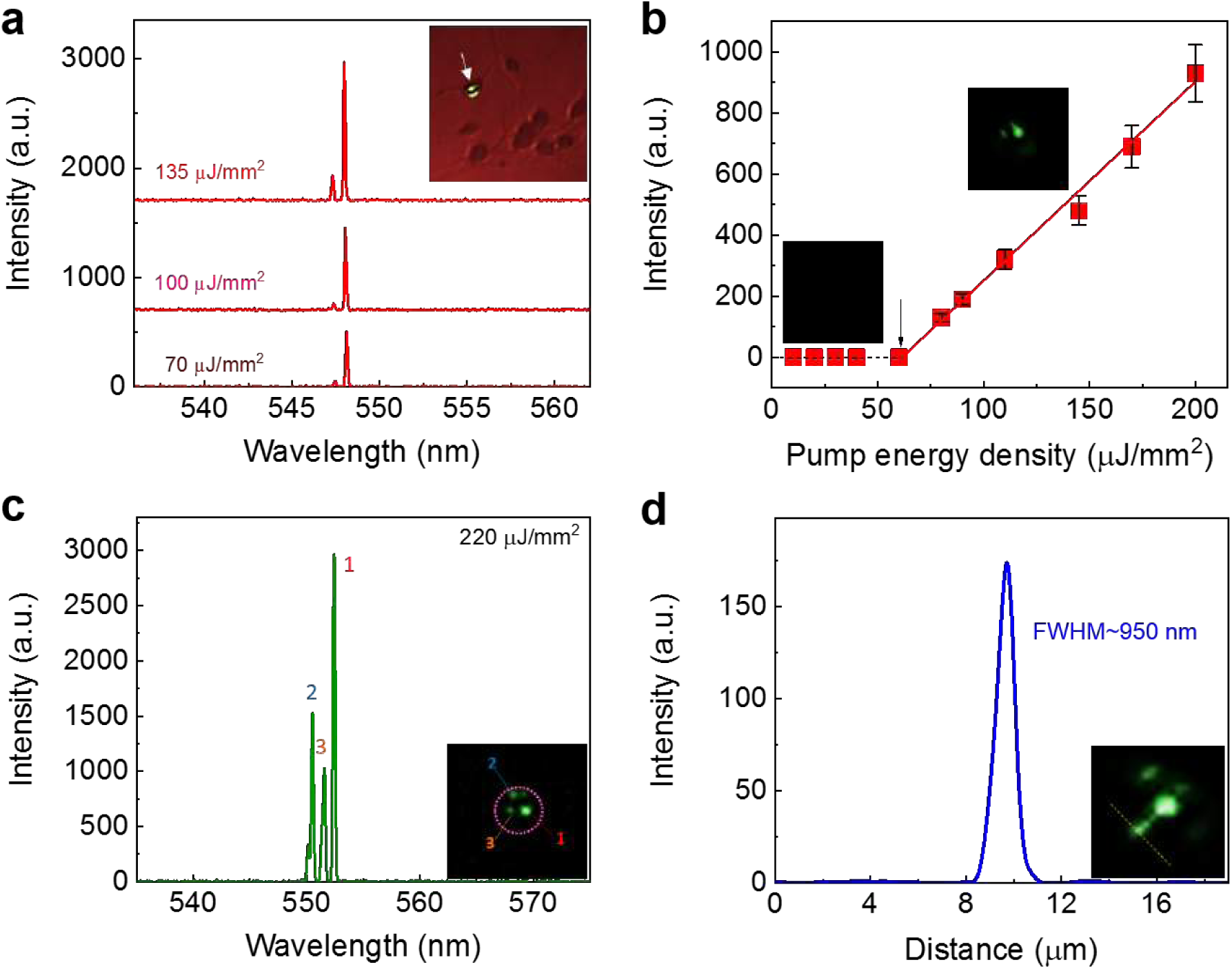
Lasing in live neurons. **(a)** Lasing spectra from a single live neuron at the resting potential under various pump energy densities. The signal-to-noise ratio for the lasing peaks is >500. The neuron was stained with 25 µM OGB-1 solution (see “Methods and materials”). The inset shows a brightfield CCD image of the neuron with subcellular laser emission in the soma, as indicated by the arrow. **(b)** Spectrally integrated (540 nm - 550 nm) laser output as a function of pump energy density extracted from (a). The solid line is the linear fit above the lasing threshold, showing a lasing threshold of 60 μJ/mm^2^ (shown by the arrow). Error bars are obtained based on the power variation of the pump laser. The inset CCD images show an example of neuron laser emission below and above the lasing threshold. **(c)** Multiple lasing peaks that correspond to the laser emissions from a single neuron that have different intracellular dye concentrations. **(d)** FWHM extracted from the dashed line in the CCD image, which gives a submicron resolution of 950 nm. The dashed line measures the lasing profile over a single dendrite.

### Laser recording of spontaneous neuron activities

Our next aim is to demonstrate optical recording/imaging of intracellular spontaneous calcium signals in neurons by using neuron laser emission. The pump energy density was fixed at 60 μJ/mm^2^. As discussed previously, at the resting potential, no lasing emission would be observed. However, with spontaneous neuronal activities, calcium spiking occurs, resulting in the emergence of laser emission. The calcium transients of an individual neuron caused by spontaneous soma activities were recorded at two different subcellular locations (site 1 and 2) over 90 seconds, as plotted in Fig. 4(a). Significantly, the relative change in lasing intensity, R, ranges from 20 to 70, which is about 100-fold improvement the traditional fluorescence-based measurement, which is less than 75% (see Supplementary Fig. 2(a)). The SNR for the laser emission image can be as large as 230, as shown in Fig. 4(b). For comparison, the SNR for the fluorescence based image is 4.1 at best. Such high R(t) and SNR result from the fact that when there is no calcium, the laser emission signal is nearly zero and the noise is simply caused by the background measurement (see, for example, the right inset in Fig. 4(b)).

**Figure 4.**
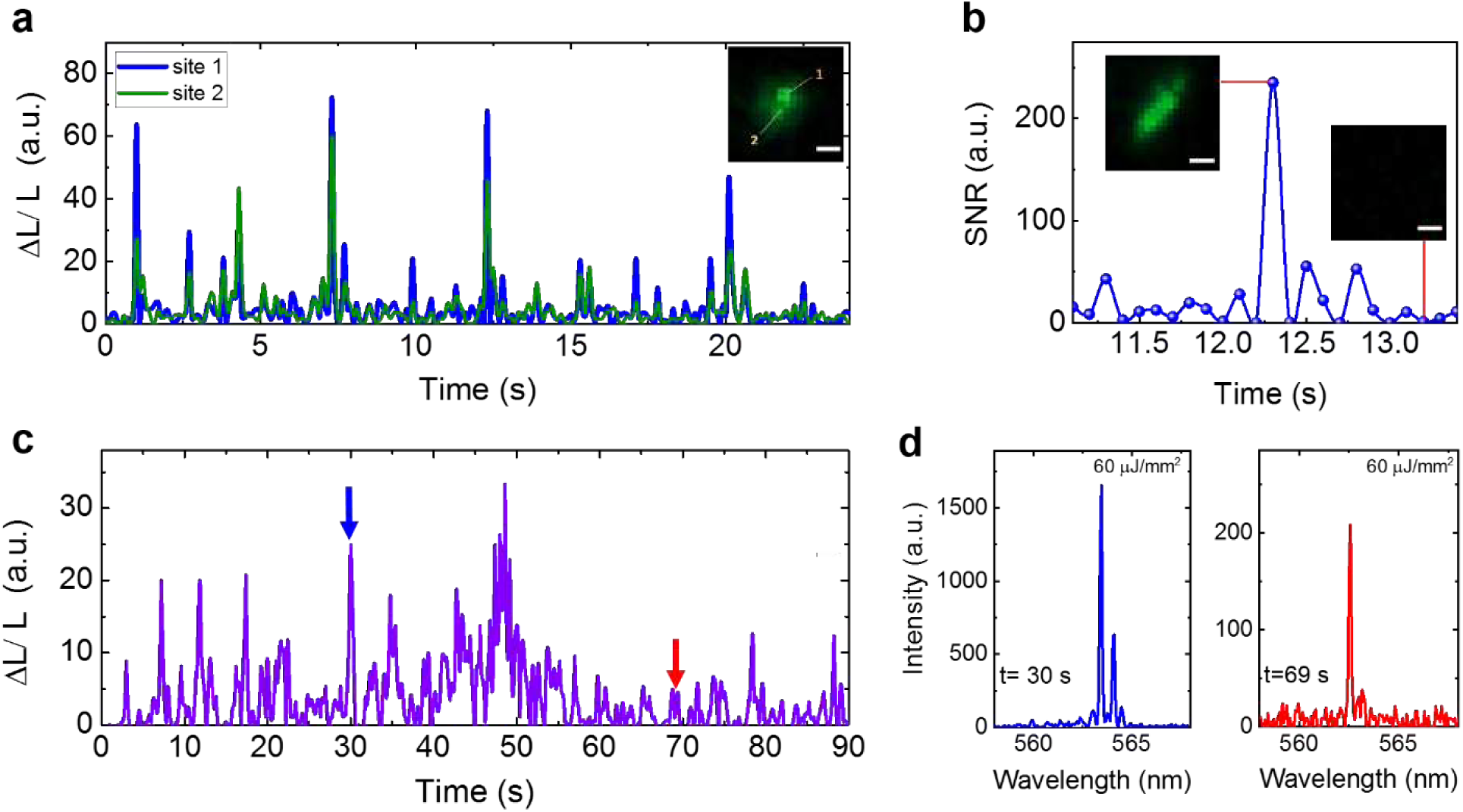
Laser recording of spontaneous spiking activities in single neurons. **(a)** Short-term laser recording of calcium transients caused by spontaneous neuronal activities over 25 seconds measured from two randomly selected sites within the same neuron. The inset shows the corresponding CCD image of laser emission from this neuron and the two subcellular sites (site 1 and 2). (b) Signal-to-noise ratio (SNR) extracted from the neuron laser recording traces from site 1 in (a). The insets show two representative images. **(c)** Long-term laser recording of calcium transients caused by spontaneous neuronal activities over 90 seconds measured from a different neuron cultured on a different mirror. **(d)** Lasing spectra measured at t = 30 s and 69 s in (c) (as marked by the two arrows), respectively. Pump energy density, 60 μJ/mm^2^. Frame rate, 10 fps. All scale bars, 2 μm.

Conventional fluorescence imaging methodologies usually consider the intracellular calcium as a whole since the fluorescence intensity appears quite uniform within the soma. Consequently, slight difference within a neuron may often be ignored. In contrast, in laser emission detection, a small difference in the gain can lead to a large difference in the laser output. In Fig. 4(a), the two sites that we selected had similar fluorescence, but produced quite distinguishable R(t) using laser recording. For example, at t = 10 s, R is approximately 22 and 8, respectively, for site 1 and 2.

Fig. 4(c) shows longer-term monitoring of the spontaneous activities within another neuron recorded by laser emission (using another FP cavity formed by another pair of mirrors). Although we used a 20 Hz pulsed laser for excitation and took images at 10 fps, we can still observe several relatively low intensity peaks following a strong peak (e.g., 11 s –15 s, 34 s – 37 s, 48 s – 52 s), which resembles the rapidly rising and then exponentially decaying waveform typically seen in the fluorescence based imaging (another example can be found at 23 s – 29 s and 33 s – 42 s in Fig. 5(c)). In the future, a high-speed CCD camera or photo-multiplier tube can be used to significantly improve the temporal resolution. In order to validate that all the CCD images recorded were from laser emission, we plot two representative spectra at 30 s and 69 s, respectively, in Fig. 4(d), in which strong laser emission can clearly be seen.

**Figure 5.**
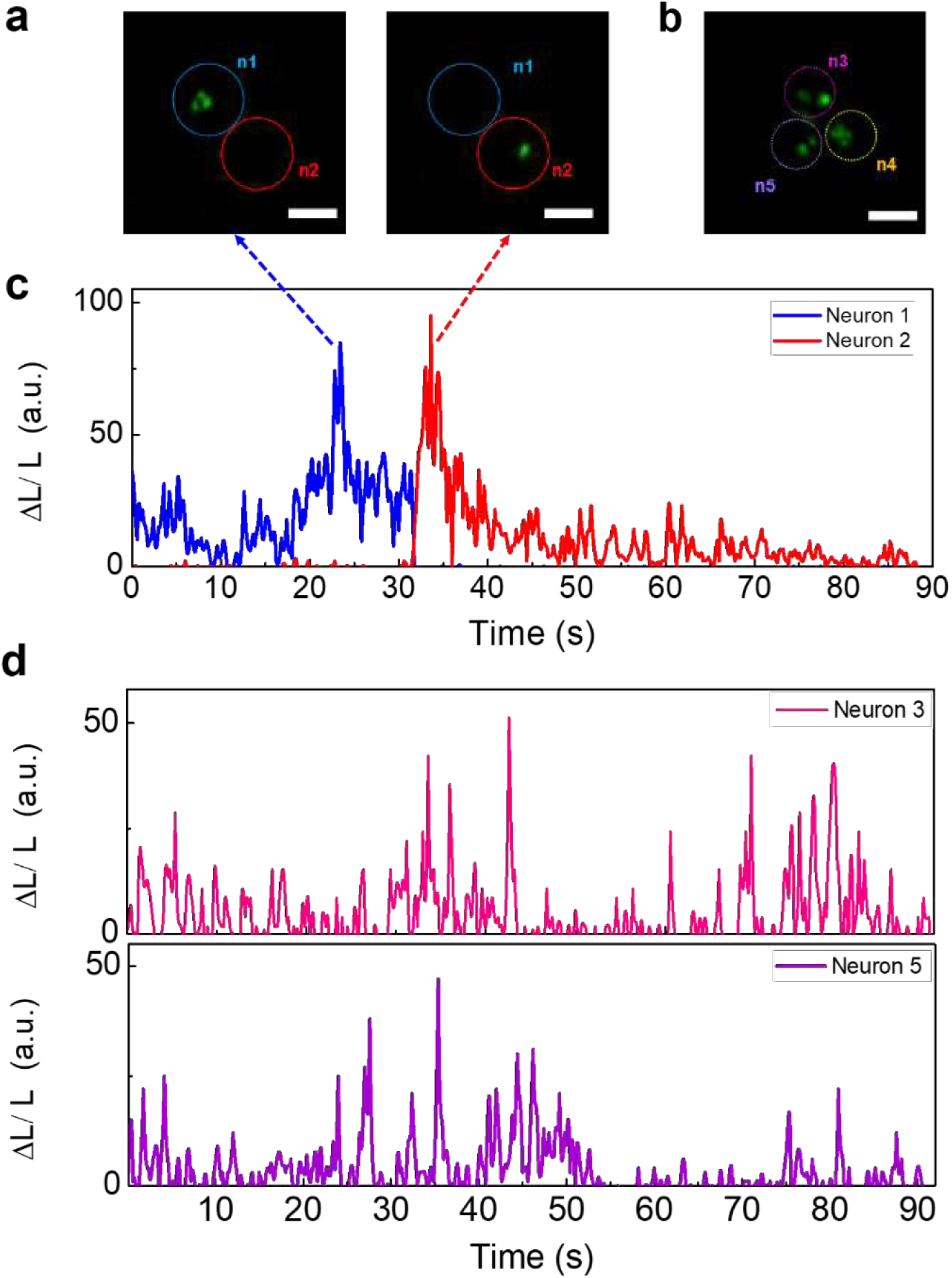
Laser recording of neuron interactions in neuronal networks. **(a)** CCD images show the laser emission switching between two neurons (n1 and n2) over time in a 2-neuron network. The left and right images were taken at t = 22 s and 32 s, respectively. **(b)** CCD image of laser emission from a neuronal network formed by 3 neurons (n1, n2, and n3). **(c)** Laser recording of calcium transients caused by spontaneous neuronal activities over 90 seconds measured from neuron-n1 and neuron-n2 in (a). **(d)** Laser recording of calcium transients caused by spontaneous neuronal activities over 90 seconds measured from neuron-n1 and neuron-n3 in (b). Pump energy density, 60 μJ/mm^2^. Frame rate, 10 fps. All scale bars, 15 µm.

In addition to single neurons, lasing from a neuron network was achieved, as demonstrated in Fig. 5. Based on the similar protocols, the calcium transients of neurons within a neuronal network were recorded and analyzed. Two types of intracellular activities, two-neuron and three-neuron networks, are presented here, as shown in the CCD images of Figs. 5(a) and (b), respectively. In Fig. 5(c), the calcium transients of neurons (n1 and n2) in neuronal networks were recorded simultaneously, in which R can be as high as 100 due to the recurring stimuli and interactions among neurons. According to the two CCD images in Fig. 5(a), we can clearly see that only n1 shows significant lasing emission at 22 s (blue trace), whereas n2 is completely dark with no lasing signal. Conversely, only n2 shows lasing emission at 32 s (red trace), while n1 turns completely dark and no lasing is observed. This may result from interconnected neurons, suggesting that neuron lasers can be biologically switched on/off based on intercellular calcium dynamics^32,33^. Next, as shown in Fig. 5(b), the CCD image clearly shows the laser emissions of three individual neurons (n3, n4, and n5). The calcium transients of neurons in neuronal networks were recorded simultaneously in Fig. 5(d).

### Laser recording of stimulated neuron activities activated by ultrasound

Finally, laser recording of stimulated neuron activities was performed, in which a low-intensity pulsed ultrasound was applied to activate neurons absorbed with piezoelectric BaTiO3 nanoparticles, which have been used previously for wireless, non-invasive, and targeted electric stimulation in neurons^34-36^. A number of studies have utilized BaTiO_3_ nanoparticles for drug delivery, imaging in *ex vivo* tissues, *in vitro* cell cultures, and *in vivo* animal or tissue models, due to their high biocompatibility^37-40^. To ensure that the BaTiO_3_ nanoparticles were well attached to the cell membrane, the BaTiO_3_ nanoparticles were first coated with a lipid layer DSPE-PEG-5000 to graft PEG onto the nanoparticles^41^. By using ultrasound as remote activation of piezo-nanotransducers, acoustic waves are transferred to electric signals for neuron stimulation, as illustrated in Fig. 6(a). Consequently, the neuron laser can be modulated by neuron’s stimulated activities, which in turn are controlled by switching on and off ultrasound.

**Figure 6.**
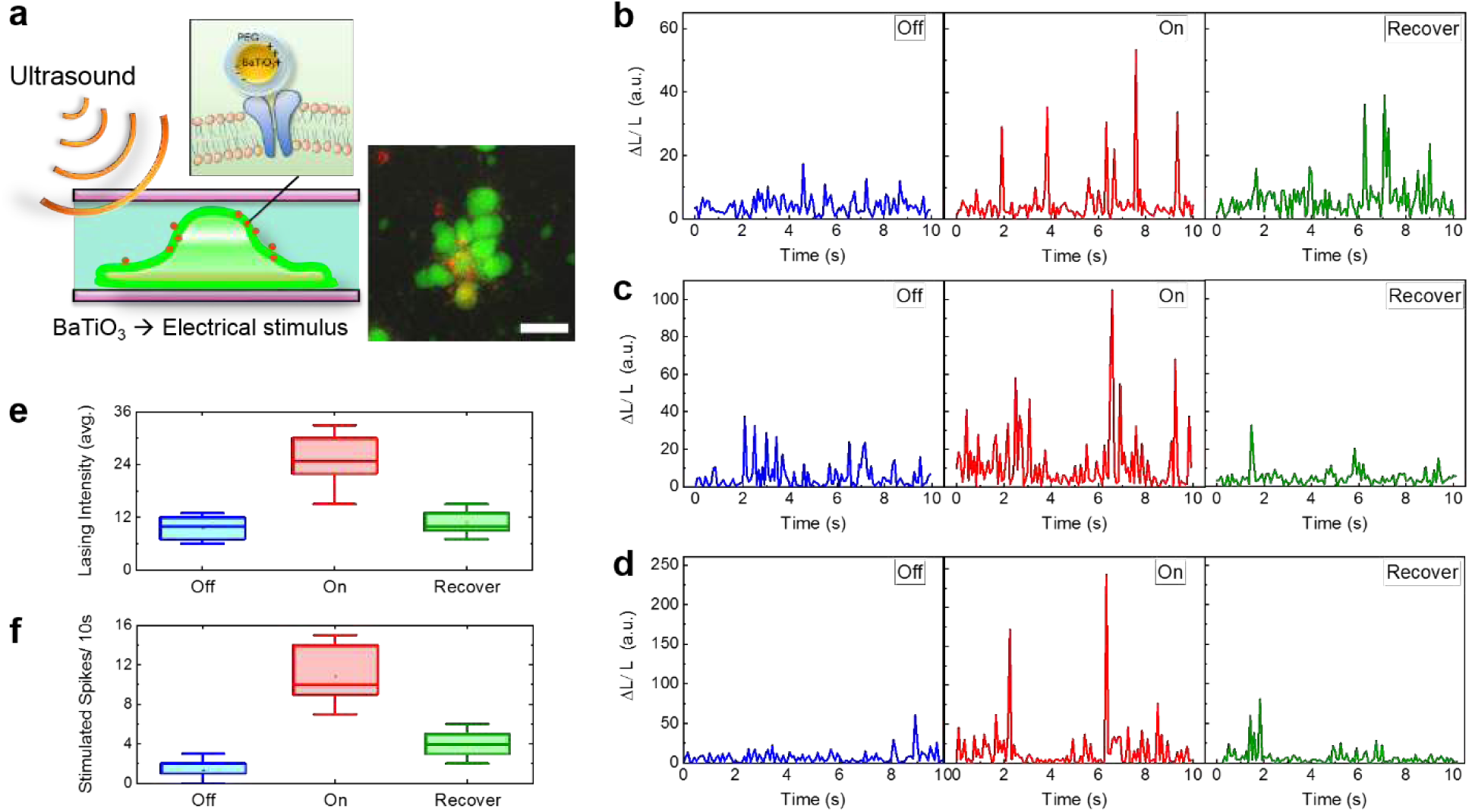
Ultrasound stimulated lasing in neuronal networks. **(a)** Schematic showing the setup of ultrasound stimulated neuron lasers. Acoustic waves (500 kHz, 2 kPa) were transformed to electrical stimuli by using piezoelectric BaTiO_3_-PEG nanoparticles adsorbed on cell membranes, as shown in the inset. The confocal microscopic image (right) shows multiple neurons (in green) with multiple BaTiO_3_-PEG nanoparticles (in red) attached. Scale bar, 20 µm. **(b-d)** Recorded stimulated spiking activities from different neuron networks before (blue curve), during (red curve), and after ultrasound stimulation (recovery) (green curve). **(e)** Corresponding statistics of the averaged lasing intensity of all effective spikes before, during, and after stimulation in 10 seconds. **(f)** Corresponding statistics of the total number of “stimulated spikes” before, during, and after stimulation in 10 seconds. Here, stimulated spikes were defined as the events when the relative change of laser intensity is above 20. The statistics was calculated based on 12 different neuron networks. Pump energy density, 60 μJ/mm^2^. Frame rate, 10 fps.

In this section, all the neurons were cultured into large neuronal networks, stained with 25 µM OGB-1 solution, and adsorbed with BaTiO_3_-PEG nanoparticles before ultrasound stimulation. In order to confirm that the neurons were activated by ultrasound mediated by BaTiO_3_ nanoparticles, we first measured the fluorescence traces of calcium signals in Supplementary Fig. 3 (by removing the top mirror). We can see that the fluorescence intensity and hence the relative change are significantly increased (by 2-4 times) for the neurons with BaTiO_3_ nanoparticles compared to those without the nanoparticles, attesting to the importance of the presence of the piezoelectric nanoparticles in neuro-stimulation. Figs. 6(b) - (d) are the examples of neuron lasing before, during, and post-ultrasound stimulation (recovery). Obviously, the relative changes in lasing intensity (ΔL/L) is found to be much higher (3-15 times) during ultrasound stimulation than prior to ultrasound stimulation for all cases. However, the lasing intensity decayed gradually as soon as the ultrasound stimulation was turned off for all three cases. The neurons’ response could be fully recovered in 5 seconds after the ultrasound was off. Such reversibility allows us to perform the laser measurements on the same cell repetitively. The statistics plotted in Fig. 6(e) show the averaged lasing intensity of all effective spikes before, during, and after ultrasound stimulation within 10 seconds. Besides lasing intensity, we calculated the number of stimulated spikes before, during, and after ultrasound stimulation. The statistics plotted in Fig. 6(f) was based on 12 individual neurons from different neuron networks. Although the number of lasing spikes does not directly equate to the number of calcium transients, the increased lasing spikes indicate higher activities of the neuron network in response to ultrasound stimulation^34,36,42^. It should be noted that the ultrasound pressure (2.0 kPa) used in this work is considered to be far below the pressure that is possible to stimulate neurons directly^34^. As a control experiment, in Supplementary Fig. 4, we recorded the laser emission of neurons under ultrasound stimulation, but without BaTiO_3_ nanoparticles, which validates that ultrasound alone does not cause any side-effect that may produce the peaks in Fig. 6 during ultrasound stimulation.

## Conclusion and discussion

To the best of our knowledge, this is the first demonstration to record subcellular neuron activities on actual neuronal networks by means of laser emission. Over 100-fold enhancement in the SNR over the signal obtained with the traditional fluorescence method allows us to study neuron spiking activities with significantly improved spatial resolution. This laser recording method, combined with the wireless neuron modulation by ultrasound, also provides a non-invasive and controllable way to study neurons under external stimuli with high sensitivities.

In the near future, a few improvement strategies will be implemented towards practical use of this technology. First, for high throughput screening/imaging of neurons, it is possible to increase the beam size (i.e., field of view) so that multiple neurons can be measured simultaneously under the same pump. Second, to improve the temporal resolution, which is currently limited by the inexpensive slow-speed CCD camera (20 fps maximally), a high speed CCD camera or a PMT can be used, along with a pulsed laser of high repetition rate (>1 kHz). Third, we note that while a very high SNR (>500) is achieved in lasing emission with a spectrometer (see, for example, Figs. 3(a) and (c)), the SNR is degraded (<230) when measured with the CCD camera. This is due to the low dynamic range of the CCD camera and the fact that the CCD collects both the narrow band lasing emission and the broad band background outside the lasing spectrum (e.g., the background below 546 nm and above 550 nm in Fig. 3(a)). A better detector with a larger dynamic range and a narrow band-pass filter can be used to further improve the SNR in the image. Finally, since the laser performance (such as lasing threshold and peak wavelength) depends on the dye concentration^16,43^, the labeling process needs to be standardized. The cavity length may also affect the lasing performance. Recently, we developed an FP cavity with SU-8 spacers of a fixed height microfabricated on the mirror, which ensures a consistent laser cavity length and highly repeatable lasing emission wavelength^44^.

As an outlook, we discuss a few possible research directions to advance the laser emission technology and explore its possible applications. First, with the established analysis in this work, it is possible to develop an optofluidic laser approach for biomolecular interaction analysis and perform G-protein coupled receptors (GPCRs)-targeted drug screen with enhanced sensitivity. Mutations in GPCRs can cause acquired and inherited diseases, and thus GPCRs are considered as one of the most powerful therapeutic targets for a broad spectrum of neurological diseases. Therefore, the design and implementation of highly sensitive GPCR assays that allows rapid screening of large compound libraries to identify novel drug candidates are crucial in early drug discovery. Microfluidics will be developed to form neuron-on-a-chip devices for applications such as monitoring of neuron-glia interactions and screening of drugs for neurological diseases. Second, the strong laser signals along with a high speed detector enable the studies of rapid neural dynamics under wide field imaging. Third, biocompatible and implantable laser probes are expected to be developed. The integration of high-performance electrodes and laser emission technology may pave a way to record brain circuits and neural activities in animals *in vivo*. This may also provide a tool for studying photonic neural networks. Finally, highly selective and remote modulation of the neuronal networks with ultrasound, in combination with the laser emission detection method, will be explored.

## Methods and materials

### Neuron preparation

In this study, primary rat cortex neurons (ThermoFisher #A1084001) were used throughout the whole experiment. Before seeding the neurons, all the mirrors were autoclaved and coated with a layer of Poly-Lysine/laminin for 30 minutes in order to achieve optimal cell adhesion. Primary neurons were then prepared and cultured on mirrors in a 6-well culture dish based on the protocol provided by ThermoFisher. All the laser measurements were conducted using the neurons 3 days after seeding.

### Materials and labeling

For intracellular calcium staining, OGB-1 (ThermoFisher #O6807) was first dissolved in Pluronic-F127/DMSO (ThermoFisher #P3000MP) at 37 °C and then diluted with phosphate buffer saline (PBS) to 100 µM. Before cell staining, 3/4 of the cell medium was first removed (cleared) from the culture dish. Next, OGB-1 was applied to the neurons and diluted with PBS again to a final staining concentration of 25 µM. After incubation at 37 °C for 3 hours, the neurons on the mirrors were rinsed with PBS to remove excessive dyes before laser measurements. In order to calibrate the labeled intracellular concentrations, we first measured the fluorescence emission intensity of different concentrations of pure OGB-1 dye mixed with 100 nM Ca^2+^ using a fluorescence microscope. Under the same experiment settings, we then measured the fluorescence emission intensity of cultured neuron cells and correlated it with the previous dye concentrations.

For ultrasound stimulated experiments, piezoelectric BaTiO_3_ nanoparticles (Sigma-Aldrich #467634-25G) were coated with 1,2-distearoyl-*sn*-glycero-3-phosphoethanolamine-*N*-[methoxy(polyethylene glycol)-5000] (DSPE-PEG, Nanocs) in order to reach a stable nanoparticle dispersion in aqueous solution. 5 mg of plain nanoparticles were dispersed in 5 mL of aqueous solution containing 1 mg/mL DSPE-PEG by using a tip sonicator (20 W for 120 s). The BaTiO_3_-PEG nanoparticles were then centrifuged (2400 rcf for 8 minutes), and the pellet was resuspended in 10 mL of ddH_2_O to obtain a 1 mg/mL of stable BaTiO_3_-PEG dispersion. Finally, 40 µL of the above solution was added to the neurons (already labeled with OGB-1 for 2 hours) and incubated for 1 hour before ultrasound stimulation.

### Optical imaging techniques

A typical confocal setup was used to excite the sample and collect emission light from the FP cavity. A pulsed OPO laser (pulse width: 5 ns, repetition rate: 20 Hz) at 475 nm was used to excite the stained neurons. The focused laser beam size was ∼30 μm in diameter. The pump energy density was adjusted by a continuously variable neutral density filter, normally in the range of 1 μJ/mm^2^ - 200 μJ/mm^2^. The emission light was collected through the same objective and then separated by a beam splitter to the spectrometer (Horiba iHR550, spectral resolution ∼0.2 nm) and a CCD (Thorlabs #DCU223C) for spectral and image analysis. The images of the laser emissions were recorded by using the CCD operated at a frame rate of 10 fps.

The fluorescence images in Fig. 1 were captured with a wide-field fluorescence microscope (by removing the top mirror in the FP cavity), whereas the corresponding lasing measurements were performed by covering the neuron samples with a top mirror. The confocal fluorescence microscopic image in Fig. 6 were taken by using a Zeiss LSM 700 confocal microscope with 488 nm and 588 nm excitation for OGB-1 and BaTiO_3_ nanoparticles, respectively.

### FP microcavities

The FP microcavity was formed by two customized dielectric mirrors (made by Evaporated Coating INC, USA). Both mirrors had a high reflectivity in the spectral range of 520 nm - 580 nm to provide optical feedback and high transmission around 475 nm for the pump light to pass through. The respective reflectivity for the top mirror and bottom mirror at the lasing wavelength (530-565 nm) was approximately 99.5%, whereas the transmission of the top mirror at pump wavelength (475 nm) was >50.0%. Microspheres (5 µm in diameter) were used as spacers to ensure the cavity length for the neuron laser. The Q-factor for the FP cavity was on the order of 10^4^ at a cavity length of 5 μm. The details of the mirror are given in Supplementary Fig. 1.

### Ultrasound stimulation setup

For ultrasound experiments, the focused ultrasound transducer was coupled through the bottom mirror via a gel while the pump laser was incident from the top mirror. The focused ultrasound transducer with 500 kHz central frequency (Olympus Inc.) and 2 kPa pressure was driven by a function generator (Stanford Research Systems, DS345) and a power amplifier (37dB, custom design) to generate US wave. A commercial calibrated hydrophone (ONDA HNC-1500) was used to measure the ultrasound distribution during the experiments.

## Data availability

Data supporting the findings of this study are available within the article and its Supplementary Information Files and from the corresponding author upon reasonable request.

## Acknowledgments

We acknowledge support from the National Science Foundation (ECCS-1607250). We would like to thank Professor Brendon Baker and William Y. Wang for the assistance of confocal microscopy and imaging.

## Author contributions

Y.C. and X.F. conceived the research. Y.C., X.L., H.Z., W.-H.W., X.T., Q.C., X.W., and X.F. designed the experiments; Y.C., X.L., H.Z. performed the experiments; W.-H.W. wrote the python code for image analysis; Y.C., X.L., W.-H.W., and X.F. analyzed data; Y.C. and X.F. wrote the paper.

## Additional information

The authors declare no competing financial interests.

