## Supplementary for "Laser Recording of Subcellular Neuron Activities"

### 1. Optical system setup for laser recording and FP cavity

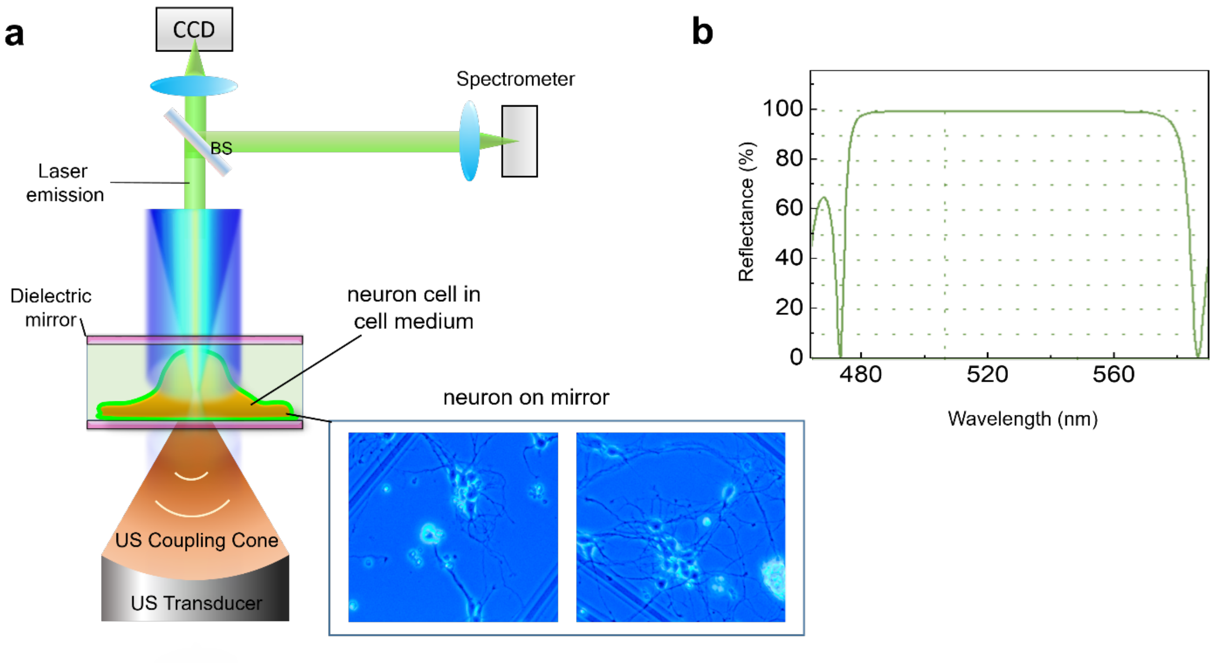


**Supplementary Figure 1. (a)** Schematic of the optical experimental setup of the laser-recording system with ultrasound stimulation. The focal beam size (spatial sampling area of the spectrum) was 30 μm in diameter. The inset figure on the bottom right shows the bright field images of neurons cultured on mirrors. The neurons were sandwiched between the two mirrors while pumped by a 475 nm pulsed excitation source. **(b)** The reflectance spectra of the top mirror and the bottom mirror. In this work the top and bottom mirror had the identical design. The reflectivity of the mirrors around the lasing emission wavelength (530-565 nm) was 99.50%.

### 2. Comparison of fluorescence recording and laser recording of neuron activities

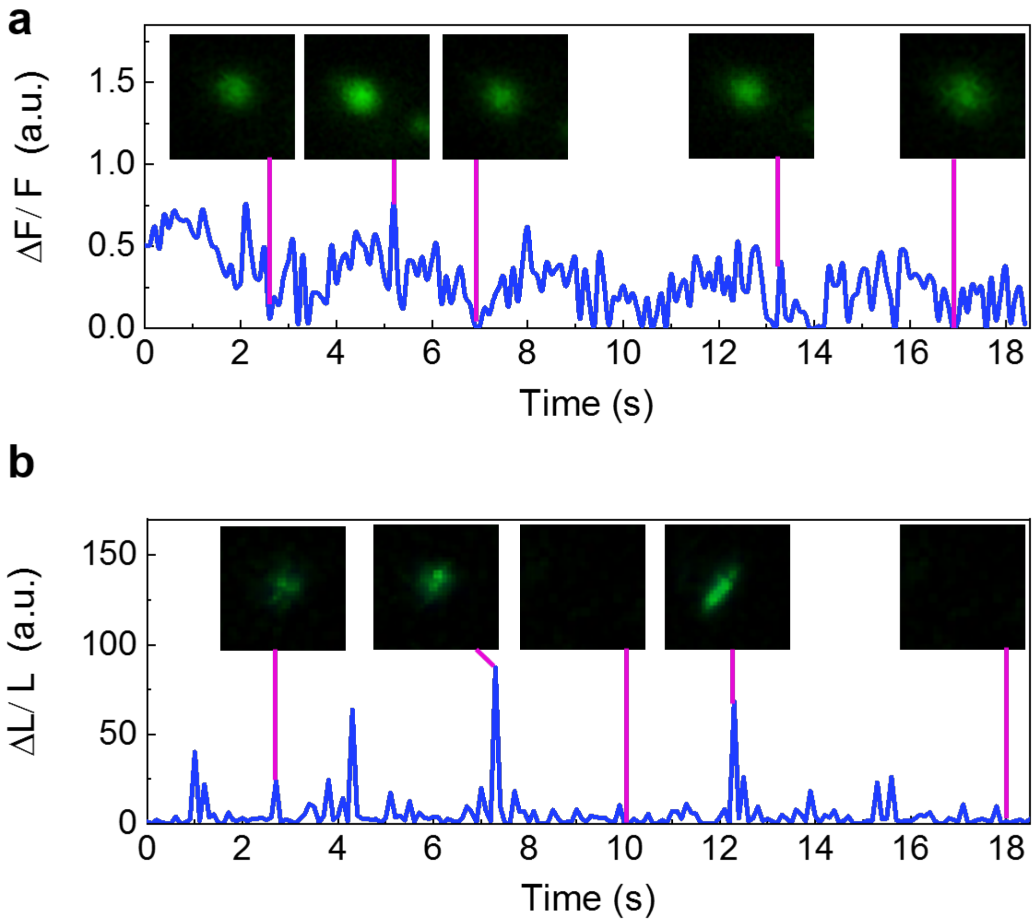


**Supplementary Figure 2. (a)** Short-term fluorescence recording of calcium transients caused by spontaneous neuronal activities over 18 seconds from a single neuron. The insets show the representative fluorescence images captured by a CCD camera at 10 fps. The SNR is less than 4.1. **(b)** Short-term laser recording of calcium transients caused by spontaneous neuronal activities over 18 seconds from a single neuron. The insets show the representative laser emission images captured by the same CCD camera at 10 fps. (a) and (b) were not measured simultaneously. The SNR is shown in Fig. 4(b). Note that the SNR for laser emission images is limited by the dynamic range of the CCD camera, which is different from the traditional images captured by scanning confocal microscopy using a photo-multiplier tube (PMT). Excitation wavelength, 475 nm. Pump energy density, 60 μJ/mm$^2$**.**

#### 3. Ultrasound stimulation effect on neurons labeled with BaTiO3 nanoparticles

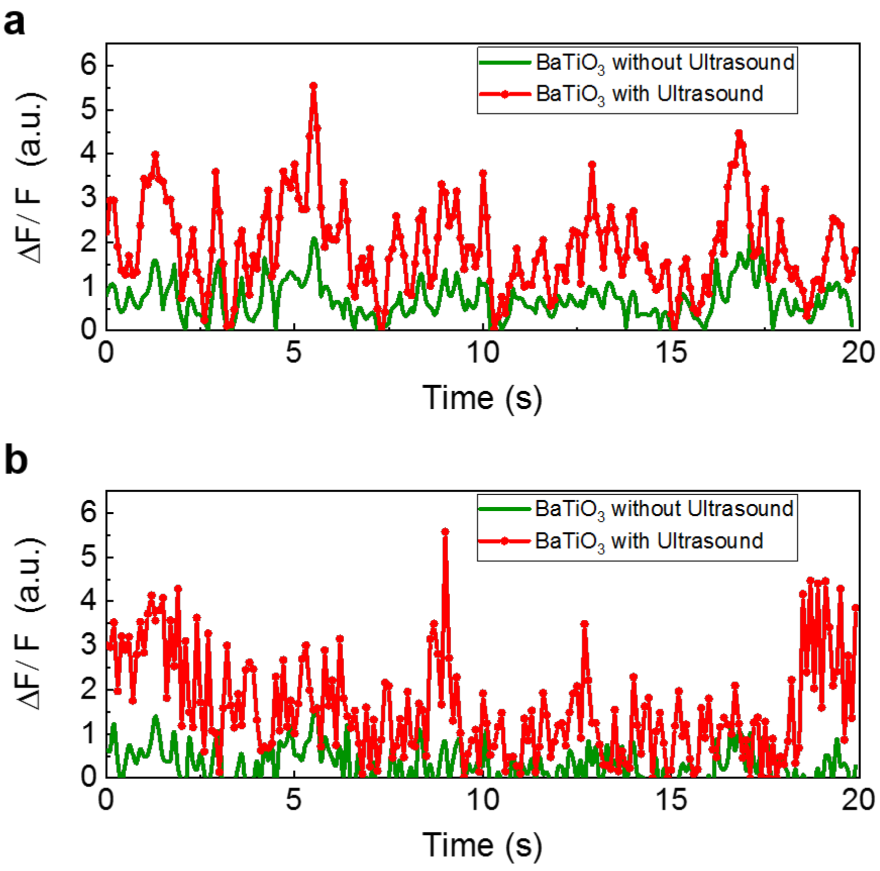


**Supplementary Figure 3. (a-b)** Fluorescence recording of two randomly selected neurons labeled with $BaTiO_3$ nanoparticles under low-power ultrasound stimulation (red traces) and without ultrasound stimulation (green traces). The relative fluorescence intensity change is significantly increased under ultrasound stimulation. Excitation wavelength, 475 nm. Pump energy density, 60 $\mu J/mm^2$.

### 4. Ultrasound stimulation effect on neuron laser

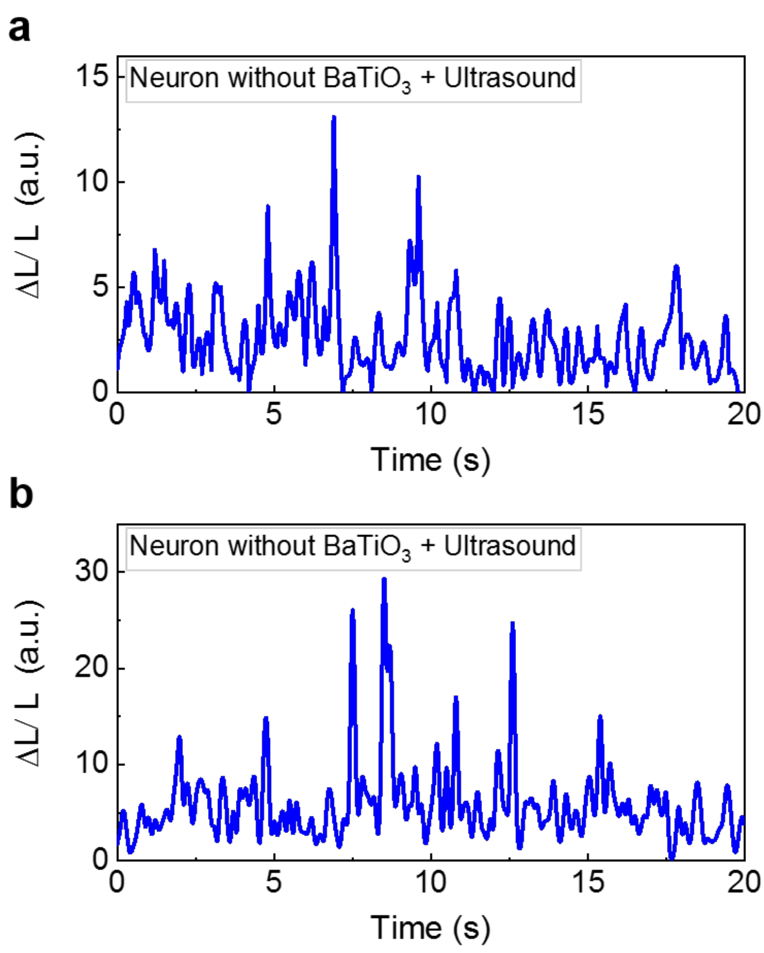


**Supplementary Figure 4. (a-b)** Laser recording of two randomly selected neurons without any $BaTiO_3$ labeling under low-power ultrasound stimulation, showing that ultrasound did not have any significant impact on neuron lasing. Excitation wavelength, 475 nm. Pump energy density, 60 μJ/mm$^2$.
